# A Hybrid Model for Predicting Pattern Recognition Receptors using Evolutionary Information

**DOI:** 10.1101/846469

**Authors:** Dilraj Kaur, Chakit Arora, Gajendra P. S. Raghava

**Affiliations:** Department of Computational Biology, Indraprastha Institute of Information Technology, Delhi, India

**Keywords:** Pattern recognition receptors, Prediction, Innate immunity, Machine learning, BLAST, Toll-like receptors

## Abstract

This study describes a method developed for predicting pattern recognition receptors (PRRs), which are an integral part of the immune system. The models developed here were trained and evaluated on the largest possible non-redundant PRRs, and non-pattern recognition receptors (Non-PRRs) obtained from PRRDB 2.0. Firstly, a similarity-based approach using BLAST was used to predict PRRs and got limited success due to a large number of no-hits. Secondly, machine learning-based models were developed using sequence composition and achieved a maximum MCC of 0.63. In addition to this, models were developed using evolutionary information in the form of PSSM composition and achieved maximum MCC value of 0.66. Finally, we developed hybrid models that combined a similarity-based approach using BLAST and machine learning-based models. Our best model, which combined BLAST and PSSM based model, achieved a maximum MCC value of 0.82 with an AUROC value of 0.95, utilizing the potential of both similarity-based search and machine learning techniques. In order to facilitate the scientific community, we also developed a web server “PRRpred” based on the best model developed in this study (http://webs.iiitd.edu.in/raghava/prrpred/).

## 2. Introduction

Pattern Recognition Receptors (PRRs) are germline-encoded proteins that are capable of sensing invading pathogens by binding to the so-called pathogen-associated molecular patterns (PAMPs) found in pathogens or by binding to the Damage-Associated Molecular Patterns (DAMPs) which are molecules released by damaged cells. This recognition of PAMPs and DAMPs by PRRs initiates a cascade of signaling processes and activates microbicidal and pro-inflammatory responses required to eliminate infectious agents, and at the same time, represents an essential link to the adaptive immune response (1). There are four significant sub-families of PRRs— Toll-like receptors (TLRs), nucleotide-binding oligomerization domain (NOD)-Leucine Rich Repeats (LRR)-containing receptors (NLR), retinoic acid-inducible gene 1 (RIG-1)-like receptors (RLR), and C-type lectin receptors (CLRs). While TLRs and CLRs are transmembrane proteins, NLRs and RLRs are cytoplasmic proteins. These PRRs play essential roles in bacterial, viral, and fungal recognition (2). Several other PRRs such as scavenger receptors, mannose receptors, and β-glucan receptors induce phagocytosis. Other secreted PRRs, such as complement receptors, collectins, ficolins, pentraxins for instances, serum amyloid and C-reactive protein, lipid transferases, peptidoglycan recognition proteins (PGRs), XA21D, etc. have also been reported in addition to the above (3).

Various studies in the past have exhibited the importance of PRRs in diseases such as autoimmune disorders (4,5), atherosclerosis, sepsis, asthma (6), heart failure (4), kidney diseases (7), bacterial meningitis, viral encephalitis, stroke, Alzheimer’s disease (AD), Parkinson’s disease (PD) (5), immunodeficiency disorders like chronic granulomatous disease (CGD) and X-linked agammaglobulinemia (XLA) (8), Cancer (9–12) etc. and thus PRRs have emerged as an important area for therapeutic research specifically in adjuvant designing (13–16). Thus, it is vital to have a deep understanding of PRR machinery and their functional roles in innate immunity. In order to facilitate the scientific community, several web resources have been developed in the past, such as InnateDB (17), IEDB (18), IIDB (19), Vaxjo (20) and VIOLIN (21). Also, an important task is an automated classification of PRRs and Non-PRRs from a vast plethora of proteins based on their structure and function. This task could be able to aid research and efficient-therapy design. Only one prediction method (22) for sub-family classification of PRRs has been developed in the past, based on data obtained from PRRDB (23). This method, however, used relaxed criteria for dataset preparation (CD hit at 90% cutoff) due to scarce data. This dataset was subsequently used for training and test of machine learning models. Since their processed dataset contains homologous sequences, the model prediction results could be biased. In order to complement and overcome the limitations of the existing method, we developed a method using the largest possible dataset, derived from PRRDB 2.0 (24) database, with standard protocols. In this study, we used protocols that divide the data into five data sets in such a way that no two proteins in two different sets have more than 40% sequence similarity, without reducing the number of sequences in the dataset (25,26). In order to understand the strength and limitation of the standard similarity based approach, we evaluate the performance of BLAST on our dataset. In the second step, we developed standard machine-learning based classification models for predicting PRRs using a wide range of descriptors like residue composition and dipeptide composition (27–30). It has been shown in the past that evolutionary information provides more information than single sequence (31,32). Thus, we developed models using evolutionary information in the form of the composition of the position-specific scoring matrix (PSSM) profile (27). Finally, we developed hybrid models that combine the strength of different approaches used in this study (32,33).

## 3. Materials and methods

### 3.1 Dataset

PRRs sequences (positive data) were obtained from the database PRRDB2.0 (24). Initially, the total PRRs taken were 2727, which were reduced to 179 unique PRRs after the removal of identical UniProt ids. The negative dataset was created by collecting random sequences from Swiss-Prot (34), which were not PRRs. The negative dataset constituted of 274 Non-PRR sequences. CD-HIT (35) with a cutoff of 40% sequence similarity was then used on both positive and negative datasets, to obtain clusters with similar sequences. 106 positive clusters and 210 negative clusters were obtained. The distribution of the sequences in the clusters is shown in Figure 1. For the positive dataset, five subsets were created from the clusters obtained by CD-HIT. All the sequences in the first cluster were assigned to the first subset and the next cluster’s sequences to the subsequent subset. We continued this process until all sequences (contained in CD-HIT generated clusters) were distributed in the five subsets. A similar process was implemented for negative dataset to create 5 negative subsets. This strategy makes sure that the subsets are dissimilar to each other (< 40% similarity between sequences in two subsets), which will be beneficial for unbiased training and test of machine learning models and selection of a better classification model. The aim of this process is to create non-redundant dataset without reducing the number of proteins from the dataset (25,26).

**Figure 1.**
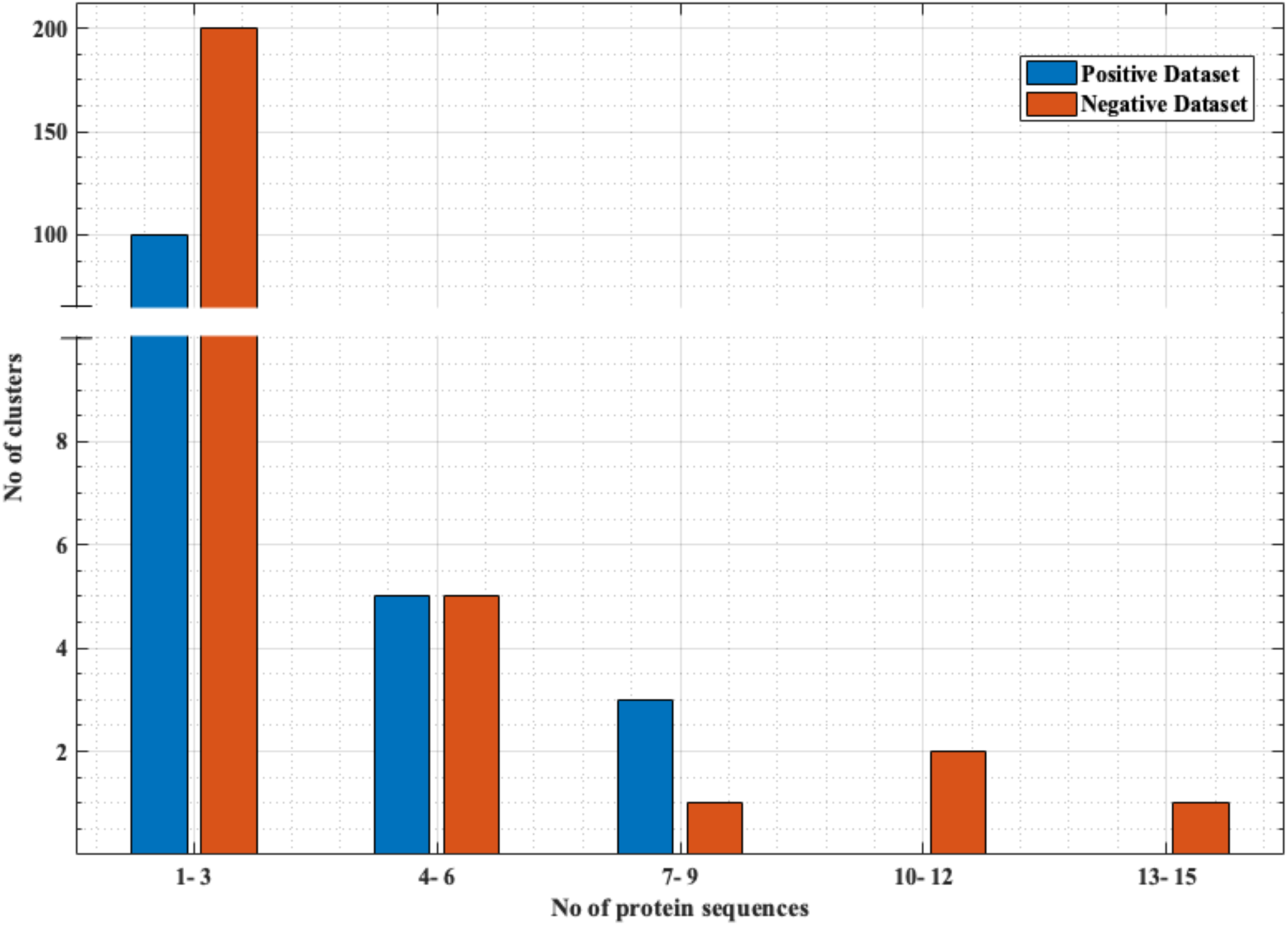
Distribution of the sequences in negative and positive clusters obtained from CD-HIT.

### 3.2 Five - fold cross validation

The performance of the modules constructed in this report was evaluated using five-fold cross-validation technique. Training and test sets were formed using positive and negative subsets. Four positive and the corresponding four negative subsets were combined to form the training set. The remaining one positive and one corresponding negative subset were combined to form the test set. This process is repeated 5 times, such that the combination of a positive subset and the corresponding negative subset is used as a test set exactly once. We employed these five training and test sets for performing five-fold cross-validation to select the best machine learning models as well as for developing BLAST similarity search based module, as explained in the next sections. Five-fold cross-validation is a standard process that has been successfully implemented in several machine learning-based studies in the past (29,36–40).

### 3.3 BLAST based similarity search

A similarity search based module was designed based on pBLAST (BLAST+ 2.7.1)(41). To evaluate the performance of this module, five-fold cross-validation was implemented. For this, a train set was used to make a local database against which the query sequences (sequences in the test set) were searched at an e-value of 0.001. The procedure is repeated five times (for each training and test set), and the evaluation metrics are noted (Results). Finally, the total positive (179 PRRs) and negative dataset (274 Non-PRRs) are combined to make a database of 453 proteins against which the user’s unseen query protein can be searched.

### 3.4 Protein Features

#### 3.4.1 Composition based features

Amino acid composition (AAC) and di-peptide composition (DPC) were obtained from Pfeature and used as features that provide residue information of a protein. AAC, for a protein sequence, is a 20 length vector where each element is the fraction of a specific type of residue in the sequence. DPC, on the other hand, is a 400 length vector that gives the composition of the amino-acids present in pairs (e.g., L-M, G-L, and so on) in the protein sequence. The detailed information can be obtained from Pfeature (42).

#### 3.4.2 Evolutionary information based features

In this study, we obtained evolutionary information for a protein using PSI-BLAST. We implemented evolutionary information in the form of PSSM-400 composition profile as a feature, similar to the previous studies (27,43–47). PSSM-400 for a protein sequence is a 20×20 dimensional vector, which is the composition of occurrences of each type of 20 amino acids corresponding to each type of amino acids in the protein sequence. For each protein sequence, PSSM matrix was created, which was then normalized and converted to a 20×20 PSSM composition vector using Pfeature’s (42) ‘Evolutionary Info’ module.

### 3.5 Machine learning techniques

We used Sci-Kit’s sklearn package, consisting of various classifiers, to develop prediction models. Each of these methods requires fixed-length feature vectors. The maximum information about proteins of variable lengths was converted into fixed vectors of equal dimensions (AAC, DPC, PSSM-400), and then these were used as input features. We used Sci-Kit’s GridSearch package to tune hyper-parameters in order to get the best performance on the training set. Subsequently, the best-learned model was used for the test. This process was implemented using five-fold cross-validation, and the average performance of five folds was evaluated. Different classifiers such as Random Forest (RF), Logistic Regression (LR), Support Vector Machine (SVM), Extra Trees (ET), K-Nearest Neighbour (KNN) and Multi-Layer Perceptron (MLP) were used to develop prediction models. All these machine learning methods have been successfully applied in many bioinformatics studies (29,36,40,48,49).

### 3.6 Performance evaluation parameters

Each model used in the study was evaluated using threshold independent and dependent scoring parameters. Threshold dependent parameters used here are Sensitivity (Sens), Specificity (Spec), Accuracy (Acc), and Matthew’s correlation coefficient (MCC). “Sens” is defined as true positive rate (TPR) i.e. correctly predicted positives with respect to actual total positives, whereas true negative rate (TNR) is defined by “Spec”. “Acc” is the ability of the model to differentiate between true positives and true negatives, while MCC is the correlation coefficient between predicted and actual classes. Following relations were used to calculate these:

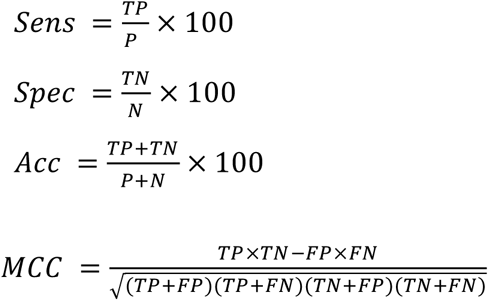

Where TP denotes correctly predicted positive, TN denotes the correct negative predictions, P denotes the total sequences in the positive set, N denotes the total sequences in the negative set, FP denotes actual negative sequences which have been wrongly predicted as positive, and FN represents wrongly predicted positive sequences. These scoring parameters are well established and have been used in many studies for model’s performance evaluation (29706944). Area under Receiver Operating Characteristic Curve (AUROC) value is a threshold independent parameter, which is calculated via the plot between True positive rate (TPR or Sens) and False positive rate (FPR or 1-Spec) (50).

### 3.7 Hybrid Models for classification

In order to improve the accuracy of the machine learning-based models further, hybrid models were constructed that combined the BLAST prediction score with the ML-based scores as done in ALGpred (51). We assigned a score of ‘+0.5’ for positive prediction (PRRs), ‘-0.5’ for negative prediction (Non-PRRs), and ‘0’ for no hits (NH). This score was added to the Machine learning-based model score. This is done for each of the sequences in the test set in a five-fold cross-validation process. Then based on this combined score, scoring metrics were evaluated for each ML model at various cutoffs.

## 4. Results

### 4.1 Prediction based on similarity search

One of the standard software which is commonly used for similarity search is BLAST. Thus we used BLAST for discriminating PRRs and Non-PRRs. In order to avoid bias, we used five-fold cross-validation, where proteins in the test set were searched against the training set using BLAST at different e-value cut-offs (Table 1). This process is repeated five times to cover all the proteins in our training sets. The positive dataset consists of 179 PRRs, and a negative dataset consists of 274 Non-PRRs. As shown in Table 1, the number of correctly predicted PRRs increased from 74.30% to 82.12% with e-value from 10-9 to 10-0 or 1. Though the performance of correctly predicted PRRs (sensitivity) increased with an increase in e-value, the rate of error (% of Non-PRRs) also increased. In the case of Non-PRRs, specificity increased from 32.48% to 49.68%, and the error rate also increased from 1.67% to 10.05% with e-value from 10-9 to 10-0. The overall accuracy of BLAST was only around 51% at e-value 10-3; due to a significant number of no-hits. This poor performance shows that BLAST is not suitable to discriminate PRRs and Non-PRRs with high precision.

**Table 1.** The performance of BLAST on training and testing dataset using five-fold cross-validation, PRRs and non-PRRs were searched at different e-values of BLAST.

| BLAST<br>(e-value) | Positive hits (Searching PRRs) |  | Negative hits (Searching Non-PRRs) |  |
| --- | --- | --- | --- | --- |
|  | PRRs (Sensitivity) | Non-PRRs (Error) | Non-PRRs (Specificity) | PRRs (Error) |
| $10^{-9}$ | 133 (74.30) | 4 (1.45) | 89 (32.48) | 3 (1.67) |
| $10^{-8}$ | 134 (74.86) | 4 (1.45) | 90 (32.84) | 4 (2.23) |
| $10^{-7}$ | 134 (74.86) | 5 (1.82) | 90 (32.84) | 4 (2.23) |
| $10^{-6}$ | 135 (75.41) | 5 (1.82) | 93 (33.94) | 4 (2.23) |
| $10^{-5}$ | 136 (75.97) | 7 (2.55) | 98 (35.76) | 5 (2.79) |
| $10^{-4}$ | 136 (75.97) | 7 (2.55) | 99 (36.13) | 6 (3.35) |
| $10^{-3}$ | **138 (77.09)** | **8 (2.92)** | **101 (36.86)** | **6 (3.35)** |
| $10^{-2}$ | 139 (77.65) | 10 (3.64) | 102 (37.22) | 6 (3.35) |
| $10^{-1}$ | 140 (78.21) | 20 (7.29) | 107 (39.05) | 7 (3.91) |
| **1** | 147 (82.12) | 65 (23.72) | 135 (49.27) | 18 (10.05) |

### 4.2 Models based on machine learning techniques

#### 4.2.1 Composition-based features

In order to develop a method for classification of PRRs and Non-PRRs, we used two main sequence composition based features viz. (i) Amino acid composition and (ii) Dipeptide composition. A wide range of machine learning techniques (e.g., SVM, KNN, RF) were used for developing prediction models. We examined the frequency of the 20 amino acids in both the positive and negative datasets. A comparison of amino acid composition between PRRs and Non-PRRs showed that residues L, N, S, and Q are more abundant in PRRs whereas A, D, E, K, and V are frequent in Non-PRRs (Figure 2).

**Figure 2.**
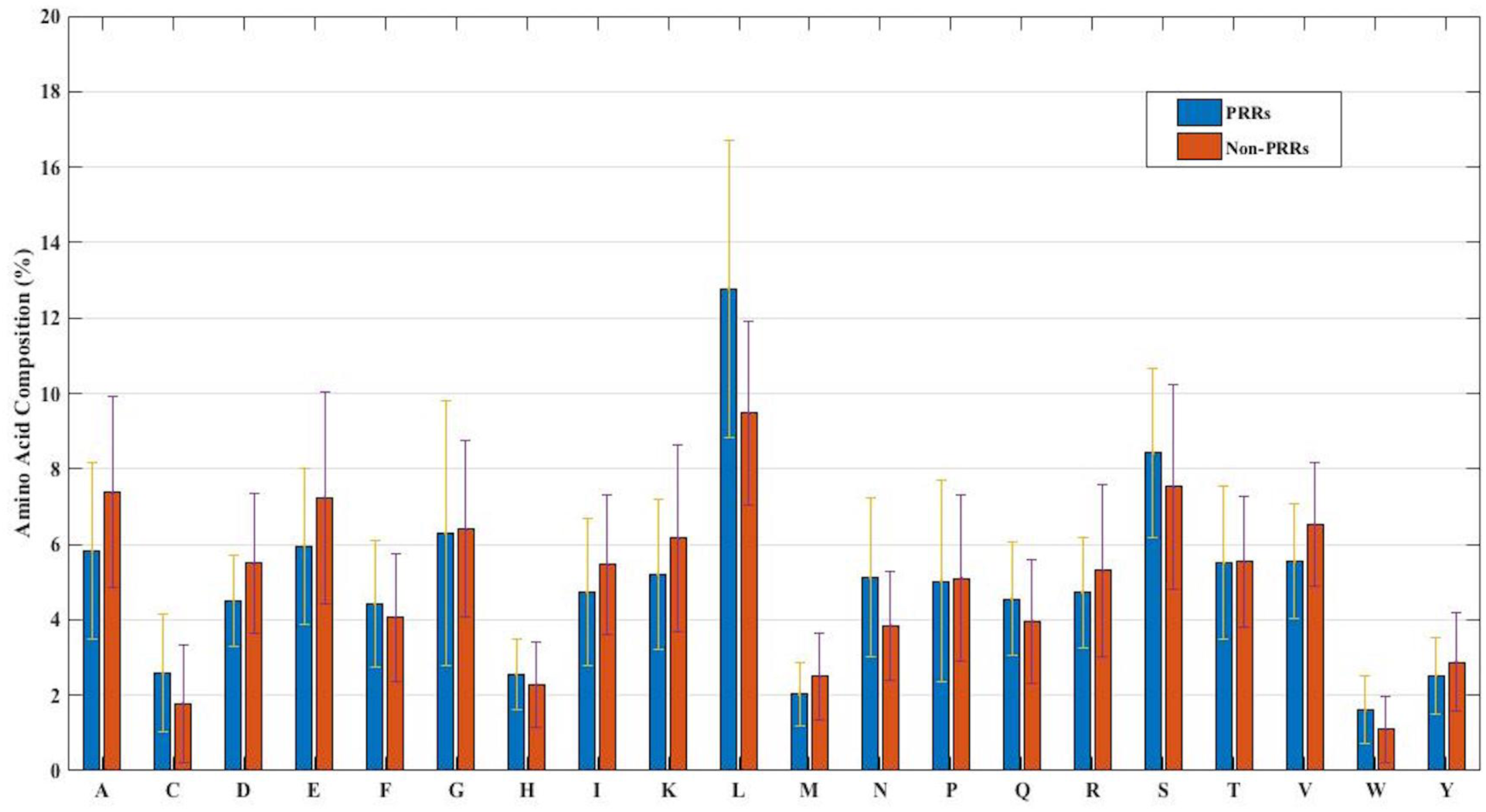
The percent amino acid composition of pattern recognition receptors and non-pattern recognition receptor proteins.

The composition of PRRs is different from the composition of Non-PRRs, as shown in Figure 2. Thus amino acid composition (AAC) feature can be used to develop models for discriminating two classes. Following machine learning techniques were used for developing binary classification models; (i) Extra-trees (ET), (ii) Random forest (RF), (iii) Support vector machine (SVM), (iv) K nearest neighbour (KNN), (v) Logistic regression (LR) and (vi) Multi-layer perceptron (MLP). As shown in Table 2, ET based models got a maximum AUROC of 0.90 with an MCC value of 0.63 on a training dataset. We achieved AUROC as 0.88 with MCC 0.63 on the test dataset.

**Table 2.** The performance of different machine learning techniques based models on PRR dataset developed using AAC of protein sequences.

| Method |  | Train Dataset |  |  |  |  | Test Dataset |  |  |  |  |
| --- | --- | --- | --- | --- | --- | --- | --- | --- | --- | --- | --- |
| Model | Hyper-parameters | Sens | Spec | Acc | AUROC | MCC | Sens | Spec | Acc | AUROC | MCC |
| ET | ne=90 | 80.71 | 82.56 | 81.73 | 0.90 | 0.63 | 77.06 | 84.08 | 82.46 | 0.88 | 0.63 |
| SVM | C=5,<br>g=0.01, k=rbf | 78.07 | 83.83 | 81.62 | 0.87 | 0.62 | 77.95 | 82.31 | 81.06 | 0.88 | 0.60 |
| RF | ne=100 | 77.82 | 81.46 | 80.08 | 0.88 | 0.59 | 77.42 | 80.85 | 79.97 | 0.87 | 0.58 |
| LR | C=1 | 77.98 | 82.50 | 80.77 | 0.86 | 0.60 | 76.12 | 81.57 | 79.57 | 0.86 | 0.58 |
| MLP | a=tanh, HL=(19,),<br>m=200, s=adam | 77.02 | 82.77 | 80.50 | 0.86 | 0.59 | 78.88 | 77.94 | 78.90 | 0.87 | 0.57 |
| KNN | al=ball_tree, nn=20,<br>w=distance | 76.17 | 79.06 | 77.91 | 0.85 | 0.55 | 77.74 | 75.00 | 76.97 | 0.86 | 0.53 |

\*g=gamma, ne=n\_estimators, k=kernel, a=activation, HL=hidden layer size, s=solver, al=algorithm, w=weight, m=max\_iter and nn=n\_neighbours.

Similarly, models were constructed using dipeptide composition and using different machine learning techniques (Table S1). The best performance was noted for LR with an average accuracy of 80.25%, MCC value of 0.59, and AUROC of 0.87 at test set, while on the training dataset, an average accuracy of 82.57% was noted with MCC value of 0.64 and AUROC value of 0.88. Overall test accuracy was 83% in the case of LR, with MCC of 0.64 and AUROC of 0.88.

#### 4.2.2 Composition of sequence profile

It has been shown in the past that the sequence profile provides more information than single sequence. Thus in this study, first, we generate sequence profile corresponding to a protein using PSI-BLAST software. In order to generate a fixed number of features, we compute the composition of sequence profile or PSSM profile (See Materials and Methods). We represent the composition of PSSM profile by PSSM-400, which has a fixed-length vector of 400 elements. We generated the PSSM-400 composition profiles for our dataset and used it as feature vectors for developing classification models. Similar to the AAC and DPC based methods, we use various classifiers such as SVM, RF, ET, MLP, etc. for training and test purposes. As shown in Table 3, our models based on evolutionary information obtained show maximum AUROC 0.87 with MCC 0.64 on the training dataset. Similarly, on the test dataset, the maximum AUROC was 0.89, with MCC 0.66. As compared to composition based prediction models, PSSM based prediction model showed higher performance in terms of MCC. In terms of AUROC, the performance of both composition and PSSM based methods was nearly the same.

#### 4.2.3 Combination of sequence and PSSM composition

The PSSM composition was combined with amino acid composition to generate a feature vector of size 420. Various classifiers were used for training and testing using five-fold cross-validation. As shown in Table 4, we got AUROC 0.89 with an MCC value of 0.66 using LR on training sets. Similarly, maximum AUROC was obtained using MLP with AUROC 0.90 and MCC 0.67 at the testing dataset. Thus, the performance has been improved as compared to using evolutionary information-based features (PSSM) or composition based features (AAC or DPC) alone. Figure 3 shows the ROC curves for different classifiers corresponding to AAC, PSSM, and the combination of AAC and PSSM.

### 4.3 Hybrid models

It is apparent from the previous results that both similarity-based approach and machine learning-based models have their own pros and cons. Thus, we made an attempt to develop a method that combines the strengths of both approaches. The e-value of 10-3 was selected for the BLAST-based similarity search method based on the hits against PRRs. Since at this e-value, the probability of correct prediction was found to be reasonably high (77.09%), and the rate of error was very low (2.92%). Though the number of no-hits was too high at this cutoff (∼80%), it was compensated by a high prediction accuracy. In order to integrate the two approaches, proteins were first classified using machine learning models. In the second step, the proteins were again classified using BLAST wherein the query proteins which showed similarity with PRRs at e-value of 10-3 were assigned as PRRs. We gave preference to BLAST over machine learning-based models in predicting PRRs due to the high probability of correct prediction of the BLAST-based similarity search method. In simple words, we used machine learning techniques for classifying proteins as PRRs and Non-PRRs when there is no BLAST hit for query protein at BLAST e-value of 10-3. This hybrid strategy improved the coverage, which was earlier missing while using BLAST alone. As shown in Table 5, the performance of machine learning techniques improved drastically when BLAST was integrated. Our best hybrid model based on PSSM achieved an accuracy of 91.39% and AUROC of 0.95 with an MCC of 0.82. The performance of all hybrid models was observed to be better than the BLAST-based similarity search and models based on machine learning techniques.

**Table 3.** The performance of different machine learning techniques-based models on PRR dataset developed using PSSM-400 of protein sequences.

| Method |  | Train Dataset (average) |  |  |  |  | Test Dataset (average) |  |  |  |  |
| --- | --- | --- | --- | --- | --- | --- | --- | --- | --- | --- | --- |
| Model | Hyper-parameters | Sens | Spec | Acc | AUROC | MCC | Sens | Spec | Acc | AUROC | MCC |
| SVM | C=10, g=0.5, k=rbf | 77.80 | 85.89 | 82.78 | 0.87 | 0.64 | 79.74 | 85.46 | 83.64 | 0.89 | 0.66 |
| LR | C=1000 | 77.31 | 86.37 | 82.84 | 0.87 | 0.64 | 80.80 | 81.07 | 81.13 | 0.89 | 0.61 |
| KNN | al=ball_tree, nn=6, w=distance | 72.80 | 83.48 | 79.36 | 0.86 | 0.57 | 78.40 | 82.50 | 81.07 | 0.87 | 0.60 |
| RF | ne=80 | 75.95 | 85.01 | 81.55 | 0.87 | 0.61 | 79.07 | 81.41 | 80.74 | 0.86 | 0.60 |
| MLP | a=logistic, HL=(14,), m=200, s=adam | 75.26 | 85.09 | 81.28 | 0.86 | 0.61 | 79.07 | 81.03 | 80.26 | 0.88 | 0.59 |
| ET | ne=70 | 80.33 | 78.79 | 79.36 | 0.88 | 0.58 | 83.73 | 74.97 | 79.15 | 0.87 | 0.59 |

\*g=gamma, ne=n\_estimators, k=kernel, a=activation, HL=hidden layer size, s=solver, al=algorithm, w=weight, m=max\_iter and nn=n\_neighbours.

**Table 4.** The performance of different machine learning techniques-based models on PRR dataset developed using the combination of composition (AAC) and evolutionary information (PSSM-400) based features for protein sequences.

| Method |  | Train Dataset (Average) |  |  |  |  | Test Dataset (Average) |  |  |  |  |
| --- | --- | --- | --- | --- | --- | --- | --- | --- | --- | --- | --- |
| Model | Hyper-parameters | Sens | Spec | Acc | AUROC | MCC | Sens | Spec | Acc | AUROC | MCC |
| MLP | a=tanh, HL=(70,), m=200, s=adam | 77.70 | 86.54 | 83.20 | 0.88 | 0.65 | 81.23 | 85.50 | 84.19 | 0.90 | 0.67 |
| LR | C=1000 | 82.97 | 83.49 | 83.34 | 0.89 | 0.66 | 83.59 | 81.49 | 82.67 | 0.90 | 0.64 |
| RF | ne=60 | 80.32 | 83.24 | 82.16 | 0.88 | 0.63 | 80.43 | 82.44 | 82.16 | 0.87 | 0.63 |
| ET | ne=100 | 77.72 | 85.25 | 82.35 | 0.89 | 0.63 | 78.96 | 83.75 | 82.13 | 0.88 | 0.63 |
| SVC | C=5, g=0.01, k=rbf | 81.65 | 83.35 | 82.73 | 0.89 | 0.65 | 80.62 | 81.56 | 81.72 | 0.88 | 0.62 |
| KNN | al=ball_tree, nn=20, w=distance | 80.20 | 76.60 | 78.12 | 0.87 | 0.56 | 80.41 | 72.88 | 76.35 | 0.86 | 0.52 |

\*g=gamma, ne=n\_estimators, k=kernel, a=activation, HL=hidden layer size, s=solver, al=algorithm, w=weight, m=max\_iter and nn=n\_neighbours.

**Table 5.** The performance of different machine learning techniques-based models on test dataset when combined with BLAST hits at e-value 10^−3^.

| Feature | Model | Hyper-parameters | Sens | Spec | Acc | AUROC | MCC |
| --- | --- | --- | --- | --- | --- | --- | --- |
| PSSM | RF | C=80 | 83.24 | 96.72 | 91.39 | 0.95 | 0.82 |
| AAC | RF | C=100 | 82.12 | 94.53 | 89.62 | 0.92 | 0.78 |
| AAC+PSSM | ET | ne =100 | 87.15 | 89.78 | 88.74 | 0.95 | 0.77 |
| DPC | SVC | C=2, g=0.01, k= rbf | 79.89 | 92.34 | 87.42 | 0.93 | 0.73 |

\*g=gamma, ne=n\_estimators and k=kernel

**Figure 3.**
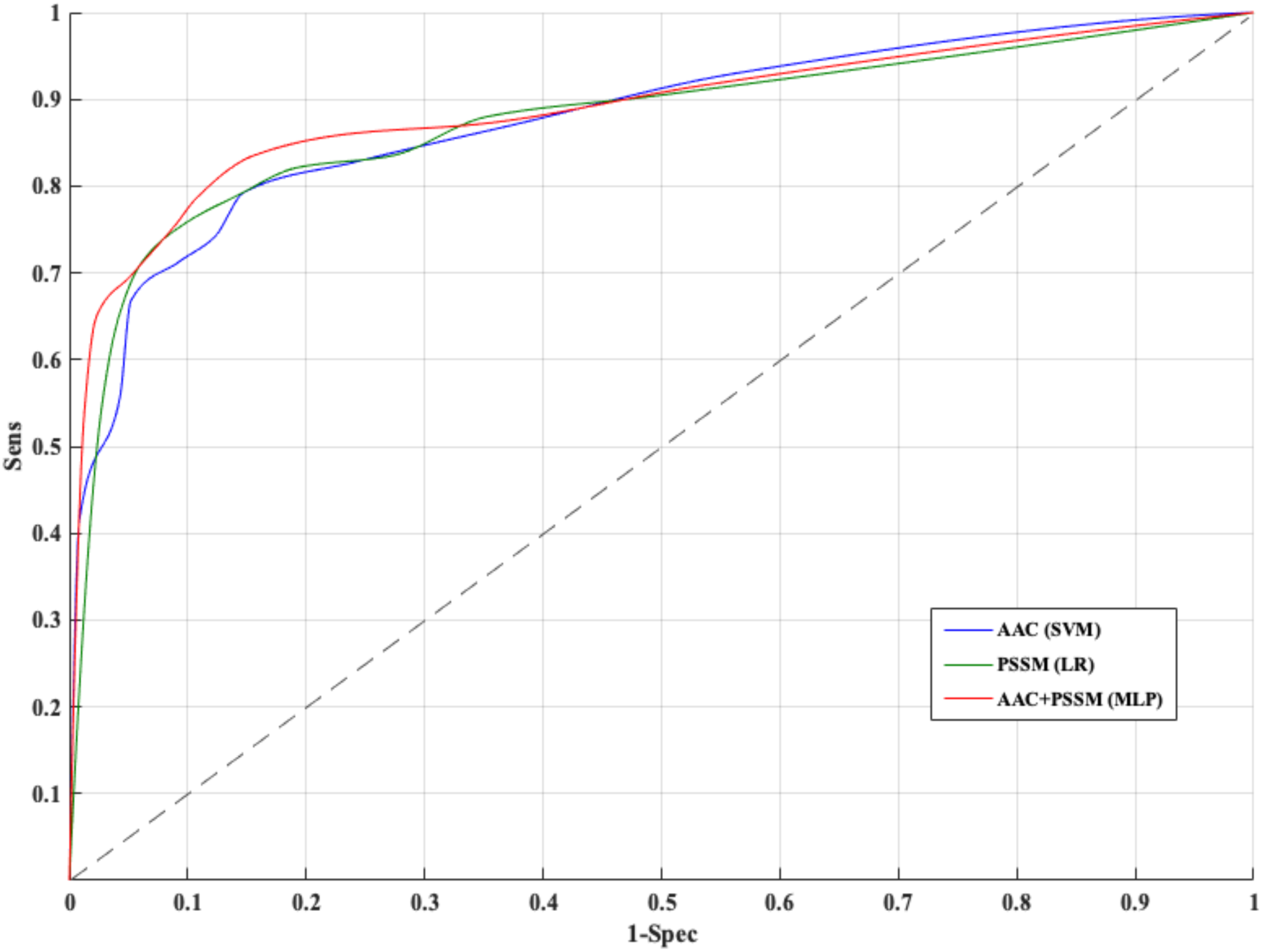
Receiver operating characteristic curves for five-fold cross-validation based on AAC, PSSM, AAC+PSSM using Support vector machine (SVM), Logistic Regression (LR) and Multi-layer perceptron (MLP) based classifier respectively.

## 5. Web-server Interface

Providing service to the scientific community is one of the primary goals of this study. We developed a user-friendly web server (http://webs.iiitd.edu.in/raghava/prrpred/), which allows users to predict whether a given protein is a pattern recognition receptor or not. The web interface of the server has two sub-modules under prediction: i) Composition Based and ii) Evolutionary Information Based. The ‘Composition Based’ module allows a user to identify a protein sequence based on Amino acid composition. This module further provides the user with the option to choose the non-hybrid method, which is only AAC based and hybrid method, which is AAC+BLAST based. The ‘Evolutionary Information Based’ module facilitates the user to predict PRRs from evolutionary information of a protein sequence. Here, the PSSM-400 composition profile for the entered protein sequence is generated and is used as a feature vector for the prediction. This module also has the facility of non-hybrid and hybrid one, which is the same as the composition based module. The web server has been designed by using a responsive HTML template for adjustment to the browsing device. Thus, our web server is compatible with a wide range of devices, including desktops, tablets, and smartphones.

## 6. Discussion

Over the past few years, there have been rapid advances in understanding innate immunity, particularly about the mechanisms by which pathogens are recognized and how the signaling molecules respond to them. Innate immunity is gaining more attention than adaptive immunity due to its role in combating the pathogens during the early stages of infection, while adaptive immunity comes later into the picture. Adaptive immunity comprises of receptors which are highly specific to antigens (52). In contrast, innate immunity consists of specialized receptors known as pattern recognition receptors (PRRs) that recognize infectious pathogens and initiate inflammatory responses for their eradication (53). Several critical implications of PRRs have been reported in the past in the context of adjuvant designing, therapeutic targets, immunomodulator design, cancer immunotherapy, etc. (52,54,55). A comprehensive database of pathogen recognizing receptors such as PRRDB (23) is highly essential to understand innate immunity. This kind of knowledge-based resources can assist researchers working in the area of innate immunity. In addition to resources, there is a need to develop methods than can annotate newly sequenced PRRs. Recently, a method has been developed using SVM for predicting PRRs and subfamilies (22). This method uses amino acid and pseudo-amino acid composition (PseAAC) for developing models using dataset derived from PRRDB (23). The prediction was based on 332 PRR sequences (containing different families) obtained from 473 sequences (that includes multiple similar UniProt IDs), which were originally present in the database, by employing CD-HIT at 90% cutoff. The model accuracy was reported to be ∼97-98%; however, such a relaxed redundancy reduction process employs sequences that can be similar up to a very high degree. In this paper, we used the dataset obtained from the recently updated version PRRDB2.0 (24) to develop classification models. The positive dataset in our case consists of PRR sequences with unique UniProt IDs, thereby first reducing the redundant data (1784 sequences) to 179 sequences. Secondly, CD hit at 40% cutoff was applied to divide both the negative (274 random Non-PRR sequences from swiss-prot) and positive datasets to five clusters each. This helped in reducing homology bias amongst the folds, and thus more precise training of the models during five-fold cross-validation.

Here, we tried various approaches to predict PRRs. We used different protein features such as composition based features (AAC and DPC) and evolutionary information-based features (PSSM) to develop machine learning-based models in order to distinguish PRRs and Non-PRRs. We also used the combination of composition based feature and evolutionary information based feature for the same. These approaches were used for the first time in the study of predicting PRRs. To do this, we used a variety of classifiers available in Sci-Kit’s sklearn such as SVM, RF, ET, MLP, etc. Firstly we tried BLAST only classification due to its simplicity and wide popularity. Though BLAST resulted in a very high accuracy (e-value of 10-3) whenever a hit was found, it was unable to predict around 80% of sequences (No-Hits) during five-fold cross-validation. Thus we employed a hybrid approach for the problem in hand, which combines ML-based methods with BLAST. The major advantage of this strategy is that the proteins which could not be predicted by BLAST alone can be predicted using ML. We tried this approach with each of the protein-features and the combination using the extensive range of classifiers. The best performance was achieved in the hybrid case of PSSM and BLAST. The formulation of this hybrid model was implemented in the free web-server. Using the web-server, for an unknown protein sequence, this model will first predict the positive (PRR) / negative (Non-PRR) class based on BLAST search against the entire database (179 PRRs+274 Non-PRRs). If the result is a ‘No-Hit’, the prediction will then be made by the RF model trained on the complete set. We believe that the work done here will be beneficial for the annotation of pattern recognition receptors and boost the ongoing research in the field of innate immunity.

