## Supplementary Table S1 for "A Hybrid Model for Predicting Pattern Recognition Receptors using Evolutionary Information"

**Supplementary File S1**

**Table S1** The performance of different machine learning techniques based models on PRR dataset developed using DPC of protein sequences.

| **Method** | | **Train Dataset** | | | | | **Test Dataset** | | | | |
| --- | --- | --- | --- | --- | --- | --- | --- | --- | --- | --- | --- |
| **Model** | **Hyper-parameters** | **Sens** | **Spec** | **Acc** | **AUROC** | **MCC** | **Sens** | **Spec** | **Acc** | **AUROC** | **MCC** |
| **LR** | C=0.1 | 78.82 | 84.91 | 82.57 | 0.88 | 0.64 | 77.46 | 81.44 | 80.25 | 0.87 | 0.59 |
| **SVM** | C=2, gamma=0.01, kernel=rbf | 77.68 | 83.52 | 81.23 | 0.89 | 0.61 | 77.02 | 80.90 | 79.62 | 0.88 | 0.58 |
| **ET** | n_estimators=80 | 72.77 | 81.32 | 78.02 | 0.86 | 0.54 | 73.03 | 79.73 | 77.22 | 0.86 | 0.52 |
| **RF** | n_estimators=60 | 73.34 | 83.05 | 79.30 | 0.86 | 0.57 | 71.23 | 77.10 | 74.95 | 0.82 | 0.48 |
| **KNN** | algorithm=ball_tree, n_neigbours =6, weight=distance | 72.40 | 77.33 | 75.31 | 0.84 | 0.49 | 67.81 | 74.40 | 72.79 | 0.81 | 0.42 |
| **MLP** | activation=logistic, hidden layer sizes,=(2,), max_iter=200,  solver=adam | 79.57 | 72.77 | 75.57 | 0.83 | 0.52 | 82.53 | 51.07 | 62.91 | 0.88 | 0.35 |

*g=gamma, ne=n_estimators, k=kernel, a=activation, HL=hidden layer size, s=solver, al=algorithm, w=weight, m=max_iter and nn=n_neighbours.
